# Contrast detection is enhanced by non-stochastic, high-frequency transcranial alternating current stimulation with triangle and sine waveform

**DOI:** 10.1101/2022.11.03.515008

**Authors:** Weronika Potok, Onno van der Groen, Sahana Sivachelvam, Marc Bächinger, Flavio Fröhlich, Laszlo B. Kish, Nicole Wenderoth

## Abstract

Stochastic Resonance (SR) describes a phenomenon where an additive noise (stochastic carrier-wave) enhances the signal transmission in a nonlinear system. In the nervous system, nonlinear properties are present from the level of single ion channels all the way to perception and appear to support the emergence of SR. For example, SR has been repeatedly demonstrated for visual detection tasks, also by adding noise directly to cortical areas via transcranial random noise stimulation (tRNS). We mathematically show that high-frequency, non-stochastic, periodic signals can yield resonance-like effects with linear transfer and infinite signal-to-noise ratio at the output. Here we tested this prediction empirically and investigated whether non-random, high-frequency, transcranial alternating current stimulation (hf-tACS) applied to visual cortex could induce resonance-like effects and enhance performance on a visual detection task. We demonstrated in 28 participants that applying 80 Hz triangular-waves or sine-waves with hf-tACS reduced visual contrast detection threshold for optimal brain stimulation intensities. The influence of hf-tACS on contrast sensitivity was equally effective to tRNS-induced modulation, demonstrating that both hf-tACS and tRNS can reduce contrast detection thresholds. Our findings suggest that a resonance-like mechanism can also emerge when non-stochastic electrical waveforms are applied via hf-tACS.

**New & Noteworthy:** Our findings extend our understanding of neuromodulation induced by noninvasive electrical stimulation. We provide first evidence showing acute online benefits of hf-tACS_triangle_ and hf-tACS_sine_ targeting the primary visual cortex (V1) on visual contrast detection in accordance with the resonance-like phenomenon. The ‘non-stochastic’ hf-tACS and ‘stochastic’ hf-tRNS are equally effective in enhancing visual contrast detection.

## 1. Introduction

### 1.1 On stochastic resonance

Stochastic resonance (SR) was discovered in the context of the hysteresis features of climate (ice age) (Benzi et al., 1981) and since then it has been generalized and studied in many systems resulting in a vast body of literature. Here we survey a few basic features of SR that are directly relevant for our paper. In general, the quality of signal transfer through a system is characterized by the following parameters at the output: amplification (or the signal strength), linearity, signal-to-noise ratio, and the phase shift.

SR is a phenomenon where the transfer of a periodic or aperiodic signal in a nonlinear system is optimized by an additive -typically Gaussian-noise (Dykman and Mcclintock, 1999). Note that originally, when SR was studied in binary systems, it represented a frequency-resonance, that is, matching the period time of the periodic signal with the mean residence time in the potential wells of the binary system driven by a stochastic carrier-wave (noise). Later the argument behind the name SR was modified to amplitude-resonance. Today, “resonance” means an optimal root-mean-square (RMS) amplitude value of the noise, i.e., amplitude-resonance at the carrier-wave RMS amplitude level for the best signal transmission.

In the initial phase of SR research, the nonlinear systems were bistable (e.g. Benzi et al., 1981). At a later stage it was discovered that monostable systems (including neurons) also offer SR (Stocks et al., 1993). Moreover, it was realized that the memory/hysteresis effects of the bistable systems actually cause a stochastic phase shift (phase noise) that negatively impacts the quality of the transferred signal (Kiss, 1996). Due to this fact, the best stochastic resonators are the memory-free Threshold Elements (TE), such as the Level Crossing Detector (LCD) (Gingl et al., 1995) and the Comparator (Stocks, 2000). The LCD device (the simplest model of a neuron) produces a short, uniform spike whenever its input voltage amplitude is crossing a given threshold level in a chosen, typically positive direction. On the other hand, the Comparator has a steady binary output where the actual value is dictated by the situation of the input voltage amplitude compared to a given threshold level: for example, in the sub-threshold case the output is “high” while in the supra-threshold case, it is “low”.

At the output of a stochastic resonator, the signal strength (SS), the signal-to-noise-ratio (SNR), the information entropy and the Shannon information channel capacity show maxima versus the intensity of the additive input noise. However, these maxima are typically located at different noise intensities. Exceptions are the SNR and information entropy which are interrelated by a monotonic function; thus they have the same location of their maxima, see the arguments relevant for neural spike trains (DeWeese and Bialek, 1995). On the other hand, the information channel capacity of SR in an LCD and in neural spike trains has the bandwidth as an extra variable controlled by the input (the higher the input noise the higher the bandwidth); thus the different location of its maximum is at higher input noise than for the maximum of the SNR (Kish et al., 2001).

It is important to note that, *in the linear response limit*, that is, when the input signal is much smaller than the RMS amplitude of the additive carrier-wave (Gaussian noise), the SNR at the output is always less than at the input, see the mathematical proof in Dykman et al. (1995). Consequently, the information content at the output is always less than at the input. On the contrary, in the nonlinear response limit, the SNR at the output can be enhanced by several orders of magnitude compared to its input value provided the signal has a small duty cycle, such as neural spikes do (Kiss, 1996; Loerincz et al., 1996). Yet, due to the unavoidable noise at the output, which is the unavoidable impact of the stochastic carrier-wave (noise), the information at the output (signal plus noise) is always less than in the *original* input signal *without the added carrier-wave* (*noise*).

Therefore, if a proper additive, high-frequency, periodic time function could be used as carrier-wave in a stochastic resonator instead of a Gaussian noise, the fidelity and the information content of the input signal could be preserved while it is passing through the nonlinear device, as we will show below. However, even in this case there is an optimal (range) for the mean-square amplitude of the carrier-wave. Thus, we call this deterministic phenomenon “*non-stochastic resonance*", which is also an amplitude-resonance where the optimal amplitude of a *non-stochastic* (instead of stochastic/noise) carrier-wave produces the optimal signal transfer via the system.

### 1.2 Non-stochastic resonance with high-frequency periodic carrier-waves

First Landa and McClintock (2000) realized that SR like phenomena could occur with high-frequency sinusoidal signals instead of noise. They successfully demonstrated their idea by computer simulations of a binary (double-well potential) SR system. Recently, Mori, et al (2022) used high-frequency, noise-free, periodic neural spikes for excitation in a neural computer model to show that SR like features on the mutual information can be achieved by tuning the frequency of these periodic excitation in the 80-120 Hz range.

Below, we show that high-frequency triangle waves can offer a noise-free signal transfer which can be exactly linear at certain conditions. Sinusoidal waves are also discussed briefly.

#### 1.2.1 The case of triangle (or sawtooth) carrier-waves, instead of noise

Earlier, in a public debate about the future of SR, one of us proposed a noise-free method by utilizing high-frequency triangle waves to improve signal transmission through threshold devices (Kish, 2007) and to reach exactly linear transfer and infinite SNR at the output. Here we summarize those arguments.

**Figure 1** shows an example of stochastic resonator hardware with additive triangle wave, as the carrier-wave, instead of noise. The same argumentation works for sawtooth wave, too. Note: the original threshold-based stochastic resonators (Gingl et al., 1995; Kiss, 1996) contain the same hardware elements where Gaussian random noise is used instead of the triangle wave. Due to the binary nature of the visual detection experiments described in this paper our focus is on sub-threshold binary signals with some additional comments about the case of analog signals.

**Figure 1.**
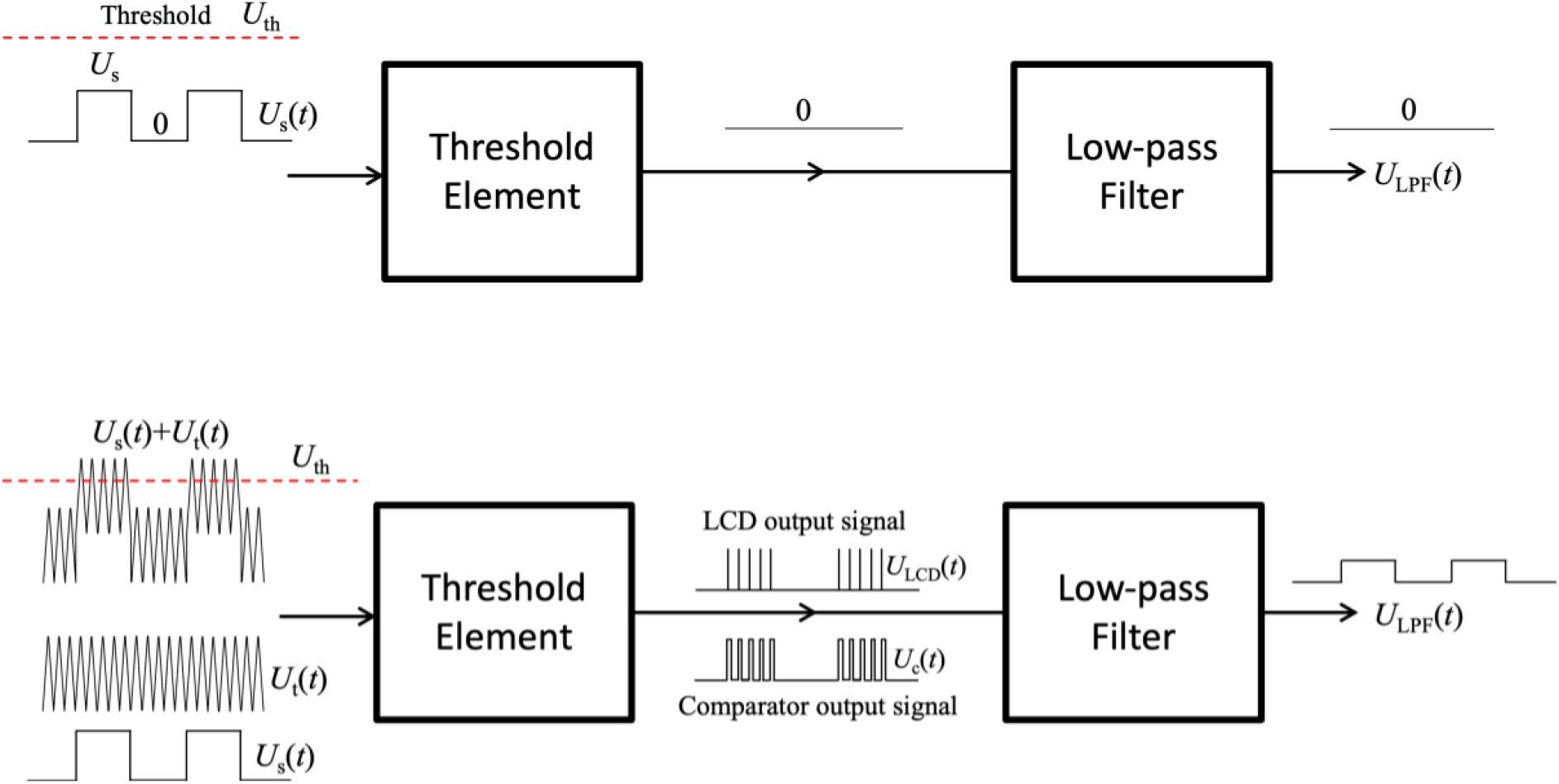
Deterministic transfer of sub-threshold binary signal through simple threshold-based stochastic resonators with a Threshold Element (TE: either a Level Crossing Detector (LCD) or a Comparator) and an additive triangle wave at the input. Note: the classical threshold-based stochastic resonators contain the same hardware elements except the triangle wave that is substituted by a Gaussian random noise. The role of the Low-pass Filter is to reduce the amount of irrelevant high-frequency products created by the carrier wave. If those irrelevant high-frequency products are not disturbing, the Low-pass Filter can be omitted. Upper part: the sub-threshold binary signal is unable to trigger the TE thus the output signal is steadily zero. Lower part: an additive, triangle wave (carrier-wave) assists the signal to reach the threshold thus it carries the binary signal over the TE. The Low-pass filter takes a short time average in order to smooth out the high-frequency components. For high-fidelity transfer, to avoid problems caused by delays or phase shifts, the frequency of the carrier-wave must be much greater than that of the binary signal. In the old stochastic resonance schemes, the carrier-wave was a noise that caused a non-deterministic component (noise) and finite SNR at the output. The new system is purely deterministic, and its SNR is infinite. Moreover, if the signal is “analog” (continuum amplitude values), the triangle wave with comparator as TE guarantees a linear transfer of the signal provided the threshold level is between the minimum and the maximum of the sum of the signal and the carrier-wave, see in (ii) below. Uth = threshold, Us = signal, Ut = noise, Ulcd = LCD output signal, Uc = comparator output signal, Ulpf = signal after low-pass filtering.

The TE is either an LCD or a Comparator. Suppose that the stable output of the LCD is zero and it produces a short uniform positive spike with height *U*_LCD_ and duration τ whenever the input level crosses the Threshold in upward direction. The Comparator’s output stays at a fixed positive value whenever the input level is greater than the threshold and stays at a lower (zero or negative) otherwise. Suppose when the input level is greater than the Threshold, *U*_th_, the Comparator output voltage *U*_c_ = *U*_H_ and otherwise it is 0. The Low-pass Filter creates a short-time moving-average in order to smooth out the high-frequency components (frequency components due to switching triggered by the carrier wave) and it keeps only the low-frequency part which is the bandwidth of the signal. The parameters, such as the frequency *f*_s_ of the signal, the frequency *f*_t_ of the triangle wave and the cut-off frequency *f*_c_ of the Low-pass Filter satisfy

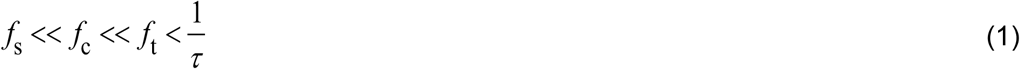

in order to transfer the signal with the highest fidelity.

The upper part of **Figure 1** shows the situation without carrier wave: the sub-threshold binary signal is unable to trigger the TE thus the output signal is steadily zero. The lower part of **Figure 1** shows the situations where an additive, triangle wave assists the signal to reach the threshold thus it carries the binary signal over the TE resulting in a nonzero output signal.

##### i) The case of Level Crossing Detector

If a constant input signal plus triangle wave can cross the threshold, the LCD produces a periodic spike sequence with the frequency of the triangle wave. In this situation, the time average of this sequence is *f*_t_τU_LCD_ therefore, for the binary input signal, the output of the LPF will be binary with amplitude values:

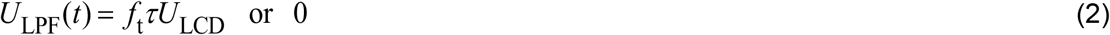

Thus, the binary input signal is restored at the output of the LPF without any stochasticity (noise) in it. The only deviation from the input signal is a potentially different amplitude (non-zero amplification) and some softening of the edges dues to the LPF depending on how well Relation 1 is satisfied.

In conclusion, with an LCD as TE, regarding the amplitude resonance versus the carrier wave amplitude *U*_t_, there are three different input amplitude ranges:

a. *U*_s_ +*U*_t_ < *U*_th_ then there is no output signal
b. *U*_th_ < *U*_s_ +*U*_t_, *U*_s_ < *U*_th_, *U*_t_ < *U*_th_ then the binary signal is restored at the output
c. *U*_th_ < *U*_t_ then the output is steadily at the high level *U*_LPF_(*t*) = *f*_t_τU_LCD_

Therefore, the binary signal can propagate to the output only in the (b) situation when it does that without any noise contribution at the output (the SNR is infinite).

##### ii) The case of Comparator

Note, this system is very different from “Stocks’s suprathreshold SR” (Stocks, 2000), where a large number of independent comparators with independent noises are used with a common signal and an adder to reach a finite SNR. For the sake of simplicity, but without limiting the generality of the argumentation, suppose that the binary signal, *U*_s_(*t*), values are 0 and *U*_s_, where *U*_s_ ≤ *U*_th_, and the maximum amplitude of the triangle signal, *U*_t_ (*t*) is *U*_t_ and its minimum value is 0. In conclusion:

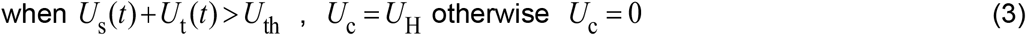

To evaluate the average output voltage of the comparator the first question is the fraction of time that the input spends over the threshold, see **Figure 2**.

**Figure 2.**
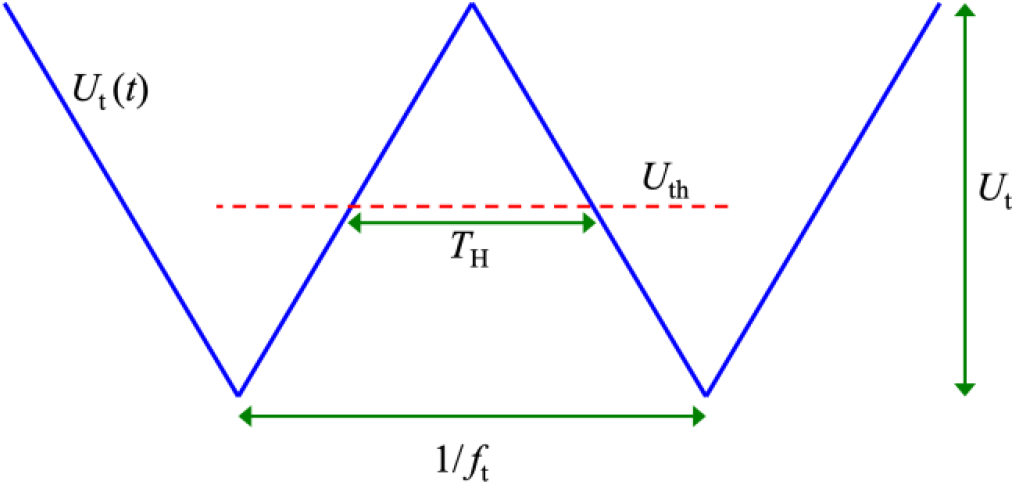
The triangle wave vs. the threshold (Uth).

This time *T*_H_ within a period of the triangle wave is the period duration 1/ *f*_t_ minus the double of the time *t*_r_ spent for rising from the minimum to the threshold:

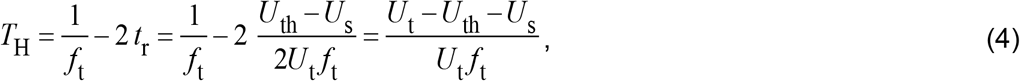

where we used that the slope *_s_* of the triangle signal with peak-to-peak amplitude is

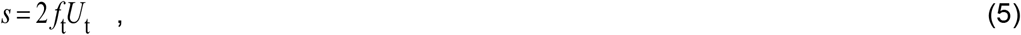

assumed that that the signal amplitude *U*_s_ is present at the input and assumed condition (3) that the signal alone is subthreshold, but the sum of the signal and the triangle wave is suprathreshold:

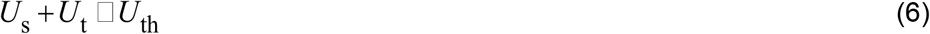

From (3) and (4), the smoothed value of the output voltage *U*_LPF_(*t*) of the LPF when the input signal amplitude is *U*_S_(*t*) = *U*_H_ :

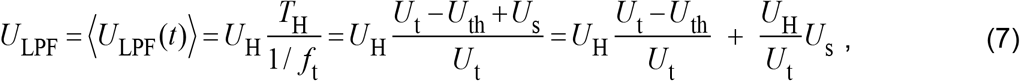

where ⟨ ⟩ denotes short-range averaged (smoothed) value discussed above.

It is obvious from the last term in the right side of Equation (7) that the signal amplitude transfers linearly through the system. Therefore, this version of our device is working distortion-free also for analog signals, not only for the present digital signal assumption.

This device is not only noise-free but also ideally linear for subthreshold signals satisfying condition (6).

In conclusion, with a Comparators as TE, regarding the amplitude resonance versus the carrier wave amplitude *U*_t_, there are two different input amplitude ranges:

a. *U*_s_ +*U*_t_ < *U*_th_ then there is no output signal
b. *U*_th_ < *U*_s_ +*U*_t_, *U*_s_ < *U*_th_, then the binary signal is restored at the output and its amplitude scales inversely with the amplitude *U*_t_ of the carrier wave. The maximal amplitude is at *U*_t_ = *U*_th_.

Therefore, the binary signal can propagate to the output only in the (b) situation when it does that without any noise (the SNR is infinite) and it has a linear transfer for analog signals.

#### 1.2.2 The case of sinusoidal carrier waves, instead of triangle waves

The above argumentations qualitatively work also for sinusoidal carrier-waves except that the linearity of the transfer is lost. The triangle carrier-wave has a Fourier series that has only odd harmonics, where the *n*-th harmonic amplitudes scale with 1/*n*^2^, that is, the strongest harmonic (the 3-rd) is 9 times weaker, and the next strongest harmonic (the 5-th) is 25 times less than the base harmonic. The qualitative difference is that the absolute value of the slope of sinusoidal carrier-wave is reduced when approaching its peak level and it is zero at the peak. The constant slope of the triangle wave is essential for the exactly linear transfer, see the mathematical proof above.

In conclusion, when sinusoidal carrier-wave is used instead of a triangle (or sawtooth) wave, the same qualitative features remain, including the zero-noise contribution at the output (infinite SNR). The exception is the linearity of transfer of analog signals via the comparator which is lost at sinusoidal carrier-wave.

### 1.3 Stochastic resonance effects on neural processing

In neural systems, it has been demonstrated that responses to externally applied stimuli were maximally enhanced when an optimal level of electrical random noise stimulation was applied. These effects were linked specifically to the opening of voltage gated sodium (Na^+^) channels in response electrical stimulation, causing a sodium influx, which in turn causes a local depolarization of the cell membrane (Onorato et al., 2016; Remedios et al., 2019; see Potok et al., 2022b for review).

In humans, early SR effects have been mainly demonstrated via behavioral signal detection tasks whereby noise was added to the periphery. For example, the detection of low-contrast visual stimuli was significantly enhanced when the stimuli were superimposed with visual noise (Simonotto et al., 1997)

Recently, similar enhancements of visual perception have been reported when noise was directly added to the cerebral cortex by the means of transcranial random noise stimulation (tRNS) in studies investigating its acute effects on visual processing (Battaglini et al., 2020, 2019; Ghin et al., 2018; Pavan et al., 2019; van der Groen et al., 2019, 2018; van der Groen and Wenderoth, 2016; see Potok et al., 2022b for review). According to the SR theory, while the optimal level of tRNS benefits performance, excessive noise is detrimental for signal processing (Pavan et al., 2019; van der Groen et al., 2018; van der Groen and Wenderoth, 2016), resulting in an inverted U-shape relationship between noise benefits and noise intensity. In consistence with SR, tRNS was shown to be particularly beneficial for visual detection performance when the visual stimuli were presented with near-threshold intensity (Battaglini et al., 2019; van der Groen et al., 2018; van der Groen and Wenderoth, 2016).

However, based on the theoretical consideration described above, a resonance-like phenomenon can be observed for non-stochastic stimulation. Here we test this prediction empirically and investigate if the response of visual cortex to around-threshold contrast stimuli could also be enhanced via high-frequency deterministic signal. We tested if triangle or sine waves can modulate signal processing in a resonance-like manner by delivering hf-tACS with triangle waveform (hf-tACS_triangle_) or sine waveform (hf-tACS_sine_) targeting the primary visual cortex (V1) of participants performing a visual contrast sensitivity task and measured their visual detection threshold. We hypothesized that resonance-like effects would be reflected in beneficial influence of high-frequency stimulation on signal processing.

## 2. Materials and methods

### 2.1 Participants

Individuals with normal or corrected-to-normal vision and without identified contraindications for participation according to established brain stimulation exclusion criteria (Rossi et al., 2009; Wassermann, 1998) were recruited in the study. All study participants provided written informed consent before the beginning of each experimental session. Upon study conclusion participants were debriefed and financially compensated for their time and effort. All research procedures were approved by the Cantonal Ethics Committee Zurich (BASEC Nr. 2018 – 01078) and were performed in accordance with the Helsinki Declaration of the World Medical Association (2013 WMA Declaration of Helsinki) and guidelines for non-invasive brain stimulation research through the COVID-19 pandemic (Bikson et al., 2020).

The required sample size was estimated using an a priori power analysis (G*Power version 3.1; Faul, Erdfelder, Lang, & Buchner, 2007) based on the effect of maximum contrast sensitivity improvement with tRNS shown by Potok et al. (2022a) (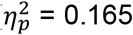, Effect size f = 0.445). It revealed that 28 participants should be included in an experiment to detect an effect with repeated measures analysis of variance (rmANOVA, 4 levels of stimulation condition), alpha = 0.05, and 90% power. We included 31 participants in experiment 1 (hf-tACS_triangle_) and 32 participants in experiment 2 (hf-tACS_sine_) to account for potential detection is potentially prone to floor effects if the contrast detected at baseline approaches the technical limits of the setup. We decided to exclude participants that are exceptionally good in the visual task and present visual contract threshold below 0.1 (Michelson contrast, see *Visual stimuli*) in the baseline, no tACS condition (for visual contrast intensity range of minimum 0 and maximum 1) in the no hf-tACS baseline condition. For outliers’ removal we used standardized interquartile range (IQR) exclusion criteria (values below Q1-1.5IQR or above Q3+1.5IQR, where Q1 and Q3 are equal to the first and third quartiles, respectively) to avoid accidental results, unlikely driven by tES, e.g., due to participants responding without paying attention to the task.

From the initially recruited sample, we excluded 7 individuals. In hf-tACS_triangle_ experiment 1: 1 participant revealed exceptional contrast threshold modulation (>Q3+1.5IQR), 1 participant had a contrast threshold below 0.1 in the baseline condition (also >Q3+1.5IQR), 1 participant stopped the session because of unpleasant skin sensations. In hf-tACS_sine_ experiment 2: 1 participant revealed exceptional contrast threshold modulation (>Q3+1.5IQR), 1 participant stopped the session because of unpleasant skin sensations, 2 participants reported frequent (75% accuracy) phosphenes sensation due to stimulation (see *Hf-tACS characteristics*).

The final sample consisted of 28 healthy volunteers (16 females, 12 males; 26.9 ± 4.7, age range: 21-39) in hf-tACS_triangle_ experiment 1, and 28 healthy volunteers (20 females, 8 males; 26.4 ± 4.4, age range: 20-39) in hf-tACS_sine_ experiment 2. Twenty of these participants completed both experimental sessions. For participants who took part in both experiments, 15 participants started with hf-tACStriangle and 5 with hf-tACSsine. The experimental sessions took place on different days with 2.6 ± 1.2 months on average apart. Delays were caused by COVID-19 pandemic (Bikson et al., 2020).

### 2.2 General Study design

To evaluate the influence of hf-tACS on visual contrast detection, we performed two experiments in which we delivered either hf-tACS_triangle_, or hf-tACS_sine_ targeting V1, during visual task performance (see **Figure 3A**). In each experiment, three hf-tACS intensities and a control no hf-tACS condition were interleaved in a random order. Our main outcome parameter in all experiments was a threshold of visual contrast detection (VCT) that was determined for each of the different hf-tACS conditions (Potok et al., 2022a). The experimental procedure to estimate VCT followed a previously used protocol to assess the influence of tRNS on contrast sensitivity (Potok et al., 2022a). In brief, VCT was estimated twice independently, in two separate blocks within each session (see **Figure 3D**). We determined the individual’s optimal hf-tACS intensity (defined as the intensity causing the lowest VCT, i.e., biggest improvement in contrast sensitivity) for each participant in the 1^st^ block of experiment 1 (ind-tACS_triangle_) and experiment 2 (ind-tACS_sine_) and retested their effects within the same experimental session on VCT data acquired in the 2^nd^ block.

**Figure 3.**
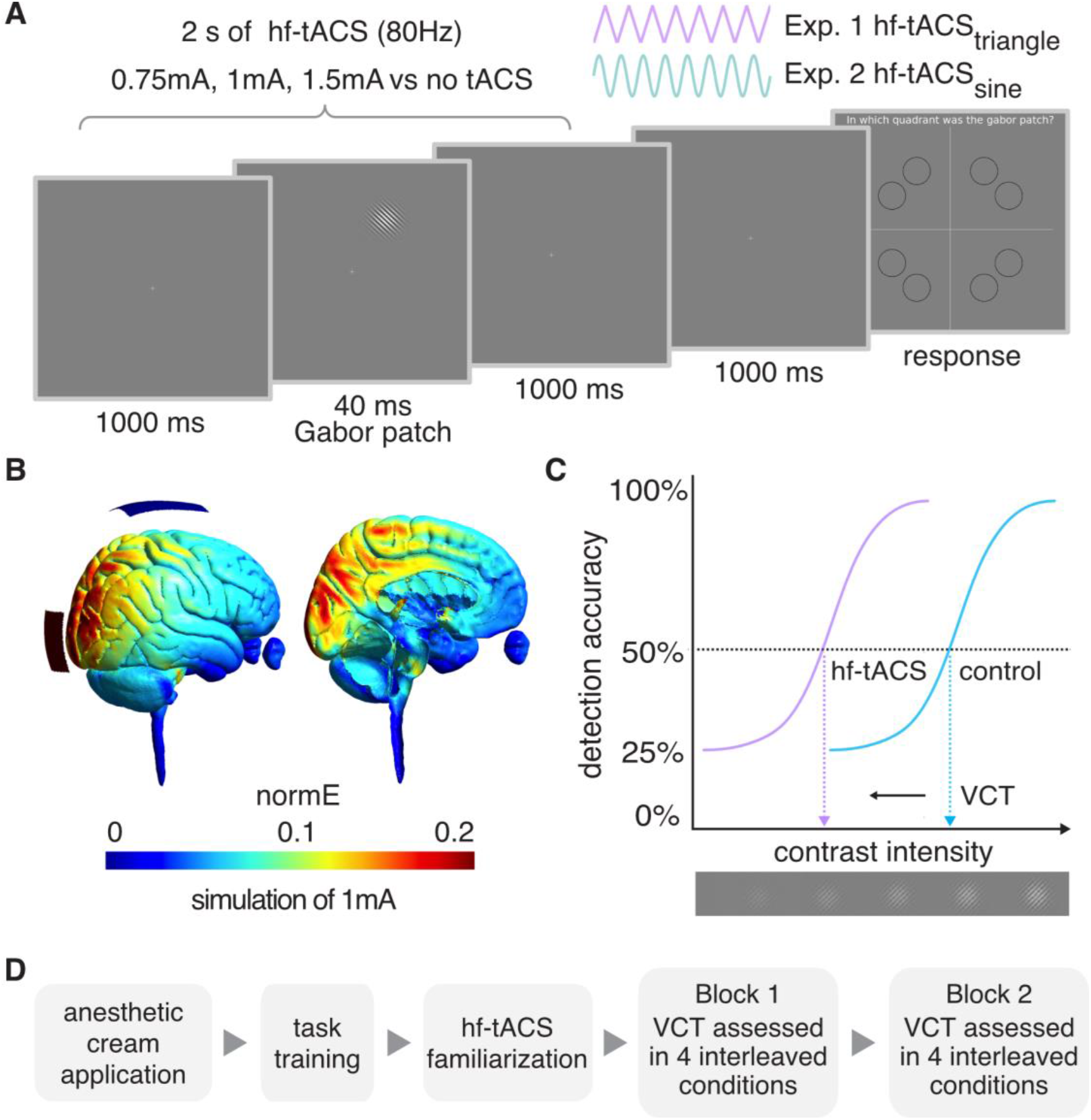
Experimental design. **A**. Example trial of 4-alternative forced choice task measuring visual contrast detection threshold (VCT). Hf-tACS was delivered for 2 s around the Gabor patch presentation. **B**. tACS electrodes montage targeting V1 and simulation of the induced electric field in the brain. **C**. Example of dose-response psychometric curves and the VCT for the 50% detection accuracy level. We hypothesize that the VCT will be lower (indicating better contrast detection performance of the participant) in one of the hf-tACS conditions (violet) than in the no hf-tACS control condition (blue). **D**. The order of measurements within each experiment. Each experimental session consisted of application of an anesthetic cream, followed by task training, familiarization protocol, and two independent VCT assessments in 4 interleaved tACS conditions (as specified in A).

#### 2.2.1 Experimental setup and visual stimuli

The experiments took place in a dark and quiet room, ensuring similar lighting conditions for all participants. Participants sat comfortably, 0.85m away from a screen, with their head supported by a chinrest. Visual stimuli were generated with Matlab (Matlab 2020a, MathWorks, Inc., Natick, USA) using the Psychophysics Toolbox extension that defines the stimulus intensity with Michelson contrast (Brainard, 1997; Kleiner et al., 2007; Pelli, 1997) and displayed on a CRT computer screen (Sony CPD-G420). The screen was characterized by a resolution of 1280 x 1024 pixels, refresh rate of 85Hz, linearized contrast, and a luminance of 35 cd/m^2^ (measured with J17 LumaColor Photometer, Tektronix™). The target visual stimuli were presented in the form of the Gabor patch – a pattern of sinusoidal luminance grating displayed within a Gaussian envelope (full width at half maximum of 2.8 cm, i.e., 1° 53’ visual angle, with 7.3 cm, i.e., 4° 55’ presentation radius from the fixation cross, staying within the central vision, i.e., <8° radius; Strasburger et al., 2011; Younis et al., 2019). The Gabor patch pattern consisted of 16 cycles with one cycle made up of one white and one black bar (grating spatial frequency of 8 c/deg). Stimuli were oriented at 45° tilted to the left from the vertical axis (see **Figure 3A**), since it was shown that tRNS enhances detection of low contrast Gabor patches especially for non-vertical stimuli of high spatial frequency (Battaglini et al., 2020).

#### 2.2.2 Four-alternative forced choice visual detection task

In both experiments, participants performed a visual four-alternative forced choice (4-AFC) visual task, designed to assess an individual VCT, separately for each stimulation condition. A 4-AFC protocol was shown to be more efficient for threshold estimation than commonly used 2-AFC (Jäkel and Wichmann, 2006). Participants were instructed to fixate their gaze on a cross in the center of the screen. In the middle of each 2.04s trial, a Gabor patch was randomly presented for 40ms in one of the 8 locations (see **Figure 3A**). A stimulus appeared in each location for the same number of times (20) within each experimental block in pseudo-randomized order to avoid a spatial detection bias. The possible locations were set on noncardinal axes, as the detection performance for stimuli presented in this way is less affected (i.e. less variable) than when stimuli are positioned on the cardinal axes (Cameron et al., 2002). Each trial was followed by 1s presentation of only fixation cross after which the ‘response screen’ appeared. Participants’ task was to decide in which quadrant of the screen the visual stimulus appeared and indicate its location on a keyboard (see **Figure 3A**). The timing of the response period was self-paced and not limited. Participants completed a short training session (10 trials), with the stimulus presented always at high contrast (0.5; for visual contrast intensity range of minimum 0 and maximum 1), in order to ensure that they understand the task (see **Figure 3D**).

VCT was estimated using the QUEST staircase procedure (Watson and Pelli, 1983), implemented in the Psychophysics Toolbox in Matlab (Brainard, 1997; Kleiner et al., 2007; Pelli, 1997). The thresholding procedure starts with a presentation of the visual stimulus displayed with 0.5 contrast intensity (Michelson contrast, for visual contrast intensity ranging 0-1; note that the stimuli were displayed for just 40ms). When participants answer correctly, QUEST lowers the presented contrast intensity. Consequently, when participants answer incorrectly QUEST increases the presented contrast. The estimated stimulus contrast is adjusted to yield 50% detection accuracy (i.e., detection threshold criterion, see **Figure 3C**). For a 4-AFC task 25% accuracy corresponds to a chance level. The remaining parameters used in the QUEST staircase procedure where set as follows: steepness of the psychometric function, beta = 3; fraction of trials on which the observer presses blindly, delta = 0.01; chance level of response, gamma = 0.25; step size of internal table grain = 0.001; intensity difference between the largest and smallest stimulus intensity, range = 1. VCT was assessed across 40 trials per stimulation condition. Four different conditions were randomly interleaved within each of 2 experimental blocks (40 trials x 4 conditions x 2 blocks; total number of 320 trials per experimental session, **Figure 3D**).

#### 2.2.3 Hf-tACS characteristics

In stimulation trials, hf-tACS (80Hz) with symmetrical triangle-(hf-tACS_triangle_) or sinewave (hf-tACS_sine_), with no offset was delivered. Stimulation started 20ms after trial onset and was maintained for 2s (see **Figure 3A**). Subsequently a screen with only fixation cross was displayed for 1 s, followed by the self-paced response time. Hf-tACS waveforms were created within Matlab (Matlab 2020a, MathWorks, Inc., Natick, USA) and sent to a battery-driven electrical stimulator (DC-Stimulator PLUS, NeuroConn GmbH, Ilmenau, Germany), operated in REMOTE mode, via a National Instruments I/O device USB-6343 X series, National Instruments, USA). The active hf-tACS conditions and no hf-tACS control condition were interleaved and presented in random order. Timing of the stimuli presentation, remote control of the tACS stimulator, and behavioral data recording were synchronized via Matlab (Matlab 2020a, MathWorks, Inc., Natick, USA) installed on a PC (HP EliteDesk 800 G1) running Windows (Windows 7, Microsoft, USA) as an operating system.

In both experiments hf-tACS stimulation (hf-tACS_triangle_ in experiment 1 or hf-tACS_sine_ in experiment 2) was delivered with 0.75mA, 1mA, and 1.5mA amplitude (peak-to-baseline), resulting in maximum current density of 60 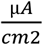 which is below the safety limits of 167 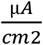 for transcranial electrical stimulation (Fertonani et al., 2015). These intensities were selected based on previous studies investigating effects of tRNS on contrast sensitivity (Potok et al., 2022a; van der Groen and Wenderoth, 2016).

Prior to electrode placement, an anesthetic cream (Emla® 5%, Aspen Pharma Schweiz GmbH, Baar, Switzerland) was applied to the intended electrodes position on the scalp to numb potential hf-tACS-induced cutaneous sensations and diminish transcutaneous effects of stimulation. To ensure that the cream got properly absorbed, it was left on the scalp for 20 min (Asamoah et al., 2019; van der Plas et al., 2020) during which participants completed task training (see *Four-alternative forced choice visual detection task* and **Figure 3D**).

To target V1 we used an electrode montage that was previously shown to be suitable for visual cortex stimulation (Herpich, 2019; Potok et al., 2022a; van der Groen and Wenderoth, 2016). The electrodes were placed on the head at least 20 min after the application of an anesthetic cream. One tACS 5×5cm rubber electrode was placed over the occipital region (3 cm above inion, Oz in the 10-20 EEG system) and one 5×7cm rubber electrode over the vertex (Cz in the 10-20 EEG system). Electroconductive gel was applied to the contact side of the rubber electrodes (NeuroConn GmbH, Ilmenau, Germany) to reduce skin impedance. The impedance between the electrodes was monitored and kept below 15 kΩ. We used electric field modelling to verify that our electrodes target V1. Simulations were run in SimNIBS 2.1 (Thielscher et al., 2015) using the average MNI brain template (see **Figure 3B**). Note, that the software enables finite-element modelling of electric field distribution of direct current stimulation without taking into account the temporal characteristics of the alternating current.

Since we used a very brief stimulation time (2 s only), fade in/out periods were not possible (Potok et al., 2021). Accordingly, some participants were able to distinguish the stimulation conditions (see *Results*). We accounted for this possible bias using a control measure and analysis of the potential transcutaneous sensations. In each session, before the start of the main experiment, participants were familiarized with hf-tACS and we assessed the detectability of potential cutaneous sensations (**Figure 3D**). The detection task consisted of 20 trials. Participants received either 2s hf-tACS (0.75, 1, and 1.5mA hf-tACS_triangle_ in experiment 1 or hf-tACS_sine_ in experiment 2) or no hf-tACS, to test whether they can distinguish between stimulation vs no stimulation. The task after each trial was to indicate on a keyboard whether they felt a sensation underneath the tACS electrodes. In experiment 2 an additional control measurement was added to assess the potential phosphene induction by the hf-tACS waveform. tACS (in lower frequency range) was previously suggested to induce visual phosphenes (Evans et al., 2021; Paulus, 2010). The protocol was the same with the only difference that this time after each trial participants indicated on a keyboard whether they perceived any visual sensations while looking on the black computer screen. The determined detection accuracy (hit rates, HR, defined as the proportion of trials in which a stimulation is present and the participant correctly responds to it) of the cutaneous sensation (experiment 1 and 2) and phosphenes (experiment 2) induced by hf-tACS served as a control to estimate whether any unspecific effects of the stimulation might have confounded the experimental outcomes (Potok et al., 2021). In the control analysis we averaged the HR for detection across tACS stimulation conditions (separately for hf-tACS_triangle_ and hf-tACS_sine_) and used the mean HR as a covariate (see *Statistical Analysis*).

### 2.3 Statistical analysis

Statistical analyses were performed in IBM SPSS Statistics version 26.0 (IBM Corp.) unless otherwise stated. All data was tested for normal distribution using Shapiro-Wilks test of normality. Partial eta-squared (small 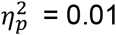, medium 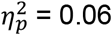, large 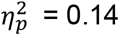; Lakens, 2013) or Cohen’s d (small d=0.20–0.49, medium d=0.50–0.80, large d > 0.80; Cohen, 1988) values are reported as a measure of effect-sizes. Variance is reported as SD in the main text and as SE in the figures. Statistical analysis of hf-tACS_triangle_ and hf-tACS_sine_ effects was analogous to the one performed to test hf-tRNS effects (Potok et al., 2022a).

#### 2.3.1 Analysis of VCT modulation in hf-tACS_triangle_ and hf-tACS_sine_ experiments

First, we tested whether baseline VCT in the no hf-tACS condition differed across the two experimental sessions using a Bayesian independent samples t-test (average baseline VCT in blocks 1-2 in experiments 1-2) using the Bayes factor (BF_10_) testing for evaluation the absence versus presence of an effect.

For all repeated measures analysis of variance (rmANOVA) models, sphericity was assessed with Mauchly’s sphericity test. The threshold for statistical significance was set at α = 0.05. Bonferroni correction for multiple comparisons was applied where appropriate (i.e., post hoc tests; preplanned comparisons of stimulation 0.75mA, 1mA and 1.5mA vs no tACS baseline).

To test the influence of hf-tACS_triangle_ on contrast sensitivity, VCT data collected in experiment 1 (hf-tACS_triangle_) were analyzed with a rmANOVA with the factors of *hf-tACS_triangle_* (no, 0.75mA, 1mA, and 1.5mA hf-tACS_triangle_) and *block* (1^st^, 2^nd^). For each individual and each block, we determined the maximal behavioral improvement, i.e., lowest VCT measured when hf-tACS_triangle_ was applied, and the associated “optimal” individual hf-tACS_triangle_ intensity (ind-tACS_triangle_). Note, that the ind-tACS_triangle_ was always selected from active stimulation conditions (i.e., even if participants performed better in the no tACS baseline, the ind-tACS intensity was defined based on the lowest VCT during stimulation). The maximal behavioral improvements in the 1^st^ and the 2^nd^ block were compared using a t-test (2-tailed) for dependent measurements. Importantly, we determined ind-tACS_triangle_ in the 1^st^ block, and then used the VCT data of the separate 2^nd^ block to test whether the associated VCT is lower compared to the no hf-tACS condition using t-tests for dependent measures. Since we had the directional hypothesis that VCT is lower for the ind-tACS_triangle_ intensity compared to no hf-tACS this test was 1-tailed. Determining ind-tACS_triangle_ and testing its effect on VCT in two separate datasets is important to not overestimate the effect of hf-tACS_triangle_ on visual detection behavior.

Similarly, VCT data collected in experiment 2 (hf-tACS_sine_) was analyzed with a rmANOVA with the factor of *hf-tACS_sine_* (no, 0.75mA, 1mA, and 1.5mA hf-tACS_sine_) and the factor *block* (1^st^, 2^nd^). Again, for each individual and each block, we determined the maximal behavioral improvement and the associated ind-tACS_sine_. We compared results obtained in the first and second block using the same statistical tests as for the experiment 1. The maximal behavioral improvements were compared using a t-test (2-tailed) for dependent measurements. We examined whether the ind-tACS_sine_ determined based on the best behavioral performance in 1^st^ block, caused VCT to be lower compared to the no hf-tACS condition when retested on the independent dataset (2^nd^ block) using t-tests (1-tailed) for dependent measures.

In both experiments to assess a general modulation of VCT induced by hf-tACS we calculated the mean change in VCT in all active hf-tACS conditions from 1^st^ and 2^nd^ blocks normalized to baseline no hf-tACS condition (hf-tACS-induced modulation).

To control for any potential unspecific effects of hf-tACS we repeated the main analyses of VCT (i.e., rmANOVA) with adding mean HR of cutaneous sensation (experiment 1, hf-tACS_triangle_ and 2, hf-tACS_sine_) and phosphene detection (experiment 2, hf-tACS_sine_) as covariate. We also tested correlations between the average HR of cutaneous sensation (experiment 1 and 2) and phosphene (experiment 2) detection and average hf-tACS-induced modulation using a Pearson correlation coefficient.

#### 2.3.2 Comparison of stimulation-induced VCT modulation in hf-tACS_triangle_, hf-tACS_sine_, and hf-tRNS experiments

We compared the effects of non-stochastic transcranial electrical stimulation (tES, i.e., hf-tACS_triangle_ and hf-tACS_sine_) and stochastic tES (i.e., hf-tRNS) on VCT. The data demonstrating the effect of hf-tRNS on VCT were taken from a previous study investigating the effects of hf-tRNS using the same behavioral paradigm (Potok et al., 2022a).

First, we tested whether baseline VCT in the no tES (no hf-tACS, no hf-tRNS) conditions differed across the three experiments using a Bayesian independent samples t-test (average baseline VCT in blocks 1-2 in hf-tACS_triangle_, hf-tACS_sine_ and hf-tRNS) using the BF_10_ testing for evaluation the absence versus presence of an effect.

Next, we tested whether a general tES-induced modulation of VCT (mean of all active stimulation conditions from two blocks normalized to baseline no stimulation condition) differed across the three experiments using a Bayesian ANOVA (tES-induced modulation in hf-tACS_triangle_, hf-tACS_sine_ and hf-tRNS experiments) using the BF_10_ testing for evaluation the absence versus presence of an effect.

Finally, we depicted tES-induced modulation of VCT as paired Cohen’s d bootstrapped sampling distributions employing an online tool (https://www.estimationstats.com; Ho et al., 2019). For each pair of control no tES (i.e., no hf-tACS in hf-tACS_triangle_, hf-tACS_sine_ and no hf-tRNS) and tES conditions (hf-tACS_triangle_, hf-tACS_sine_, hf-tRNS) a two-sided permutation t-tests were conducted. 5000 bootstrap samples were taken. The confidence interval was bias-corrected and accelerated. The reported P values are the likelihoods of observing the effect sizes, if the null hypothesis of zero difference is true. For each permutation P value, 5000 reshuffles of the control and test labels were performed.

## 3. Results

We first tested whether VCT measured during the no hf-tACS conditions differed between the experiments (i.e., average baseline VCT in hf-tACS_triangle_ and hf-tACS_sine_ experiments, see **Figure 4**). Bayesian independent samples t-test revealed that the baseline VCT measured in the no hf-tACS condition did not differ between experiments (BF_10_ = 0.29, i.e., moderate evidence for the H_0_).

**Figure 4.**
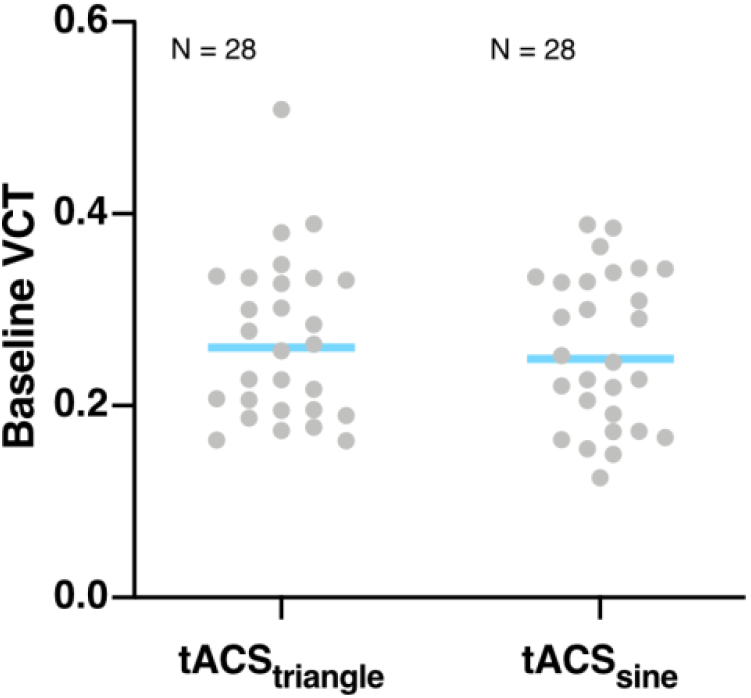
Average baseline VCT measured in the no hf-tACS conditions in hf-tACStriangle and hf-tACSsine experiments. VCT was assessed for stimuli presented with contrast intensity ranging from 0 to 1. Blue lines indicate mean, gray dots indicate single subject data.

### 3.1 Hf-tACS_triangle_ over V1 modulates visual contrast threshold

In the first experiment, we investigated whether hf-tACS_triangle_ modulates the visual contrast detection when applied to V1. We measured VCT during hf-tACS_triangle_ at intensities of 0.75, 1, to 1.5mA peak-to-baseline versus no hf-tACS control condition. We found a general decrease in VCT (F_(3, 81)_ = 3.41, p = 0.021, 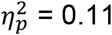) reflecting improved contrast sensitivity during hf-tACS_triangle_ (**Figure 5A**). Post hoc comparisons revealed that 0.75mA and 1mA stimulation were most effective in boosting contrast processing at a group level, which differed significantly from the no hf-tACS control condition (p = 0.033, mean difference, MD = −6.3 ± 11.62% and p = 0.024, MD = −6.33 ± 10.45%, respectively). Neither the main effect of *block* (F_(1, 27)_ = 2.43, p = 0.13) nor *hf-tACS_triangle_*block* interaction (F_(3, 81)_ = 1.6, p = 0.195) reached significance.

**Figure 5.**
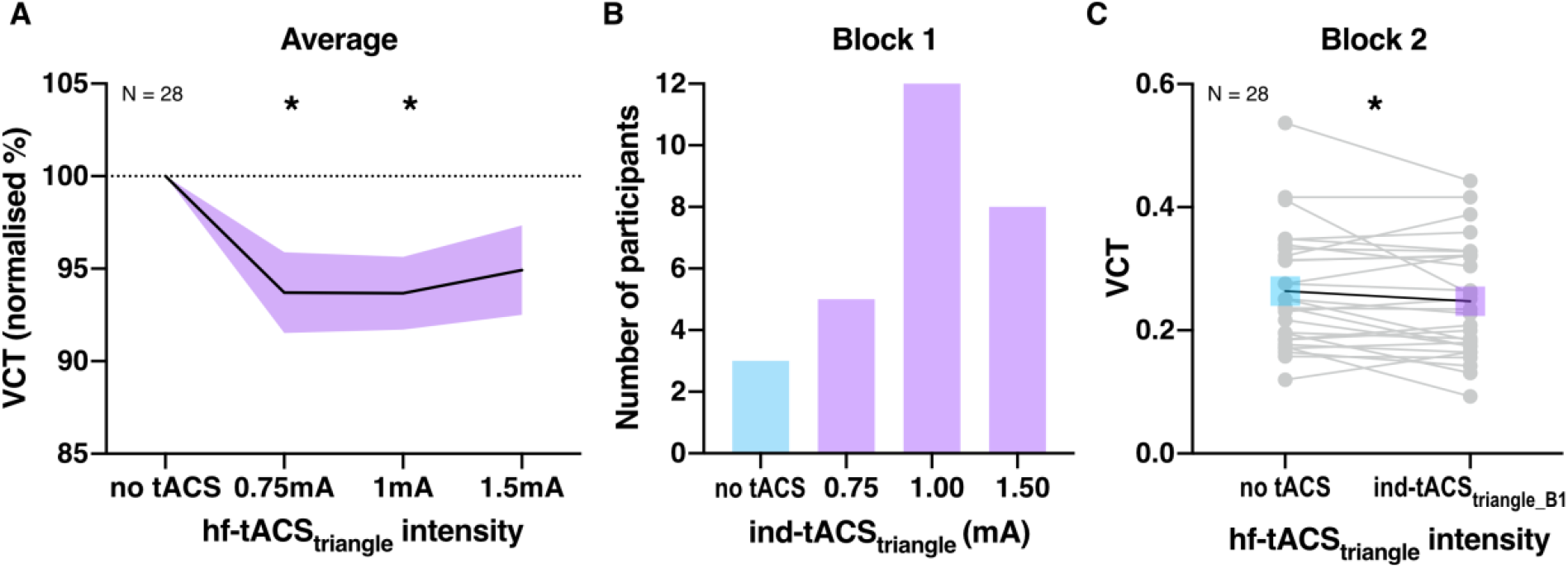
The effect of hf-tACStriangle on VCT measured in experiment 1. VCT was assessed for stimuli presented with contrast intensity ranging from 0 to 1. **A**. Effect of tACStriangle on VCT on a group level measured across 1^st^ and 2^nd^ blocks. Decrease in VCT reflects improvement of visual contrast sensitivity. VCT in hf-tACStriangle conditions normalized to the no stimulation baseline. All data mean ± SE. **B**. Individually defined optimal hf-tACStriangle based on behavioral performance during the 1^st^ block. **C**. Detection improvement effects of individualized hf-tACStriangle (selected based on block 1) measured on the independent VCT data of block 2. Gray dots indicate single subject data; *p < 0.05.

When comparing hf-tACS_triangle_-induced effects between the 1^st^ and 2^nd^ block we found that the maximal behavioral improvement (i.e., maximal hf-tACS_triangle_-induced lowering of the VCT relative to the no hf-tACS condition) were not significantly different between the 1^st^ (MD = −14.64 ± 12.6%, VCT decrease in 25 out of 28 individuals) and the 2^nd^ block (MD = −15.75 ± 15.73%, VCT decrease in 24 out of 28 individuals; t_(27)_ = 0.604, p = 0.551), additionally showing that no time effects arose from the first to the second block of measurement.

Next, we defined the optimal ind-tACS_triangle_ for each participant and examined whether its effects can be reproduced. We observed that the ind-tACS_triangle_ determined in 1^st^ block (**Figure 5B**) caused decrease in VCT compared to the no hf-tACS condition when retested within the same experimental session (t_(27)_ = 1.84, p = 0.039, VCT decrease in 18 out of 28 individuals, MD = −5.26 ± 18.23%, **Figure 5C**). Note, that the above analysis does not contain an element of intrinsic circularity because the ind-tACS_triangle_ and the VCT measure were based on independent data sets.

The cutaneous sensation control experiment revealed that some of our participants could detect hf-tACS_triangle_ conditions (HR at 0.75mA = 12.5 ± 25%, 1mA = 18.75 ± 27.74%, 1.5mA = 41.07 ± 43.68%, mean HR = 24.11 ± 27.34%). We reanalyzed our main outcome parameter by adding mean sensation detection HR as a covariate (HRs were z-scored because of non-normal distribution). The main effect of *hf-tACS_triangle_* remained significant (F_(3, 78)_ = 3.4, p = 0.022, 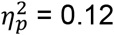). Moreover, the mean HR of cutaneous sensation detection did not correlate with the average hf-tACS_triangle_-induced VCT modulation (r = 0.181, p = 0.357), making it unlikely that transcutaneous sensation was the main driver of our results.

### 3.2 Hf-tACS_sine_ over V1 modulates visual contrast threshold

In the second experiment, we explored the effects of hf-tACS_sine_ applied over V1 on visual contrast detection. VCT was measured during hf-tACS_sine_ at intensities of 0.75, 1, to 1.5mA peak-to-baseline versus no hf-tACS control condition. We observed a general decrease in VCT with increasing hf-tACS_sine_ intensity (F_(3, 81)_ = 4.78, p = 0.004, 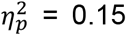) reflecting improved contrast sensitivity during hf-tACS_sine_. Post hoc comparisons revealed that the 1mA and 1.5mA stimulation were most effective in enhancing contrast processing, which differed significantly from the no hf-tACS control condition (p = 0.042, MD = −8.04 ± 13.82% and p = 0.008, MD = −6.52 ± 12.66%, respectively, **Figure 6A**). There was no main effect of *block* (F_(1, 27)_ = 0.02, p = 0.878) or *hf-tACS_sine_*block* interaction (F_(3, 81)_ = 0.5, p = 0.684).

**Figure 6.**
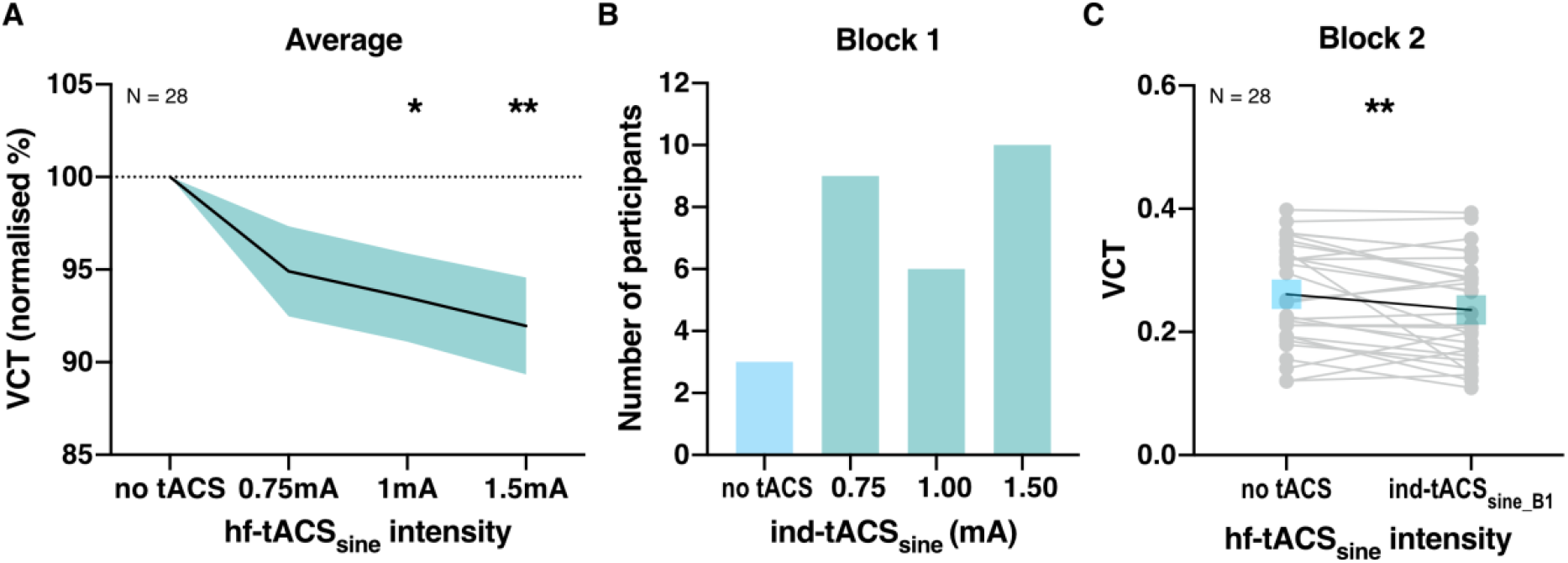
The effect of hf-tACSsine on VCT measured in experiment 2. VCT was assessed for stimuli presented with contrast intensity ranging from 0 to 1. **A**. Effect of tACSsine on VCT on a group level measured across 1^st^ and 2^nd^ blocks. Decrease in VCT reflects improvement of visual contrast sensitivity. VCT in hf-tACSsine conditions normalized to the no stimulation baseline. All data mean ± SE. **B**. Individually defined optimal hf-tACSsine based on behavioral performance during the 1^st^ block. **C**. Detection improvement effects of individualized hf-tACSsine (selected based on block 1) measured on the independent VCT data of block 2. Gray dots indicate single subject data; *p < 0.05, **p < 0.01.

When comparing hf-tACS_sine_-induced effects between the 1^st^ and 2^nd^ block we found that the maximal behavioral improvement, defined as maximal hf-tACS_sine_ induced lowering of the VCT were not different between the 1^st^ (MD = −17.78 ± 15.82%, VCT decrease in 25 out of 28 individuals) and the 2^nd^ block (MD = −18.37 ± 16.67%, VCT decrease in 22 out of 28 individuals; t_(27)_ = 0.95, p = 0.353).

We determined the optimal ind-tACS_sine_ and tested whether its effects can be reproduced. Similar to ind-tACStriangle in experiment 1, the optimal ind-tACSsine determined in 1^st^ block (**Figure 6B**) significantly lowered the VCT compared to the no hf-tACS condition when retested on the independent VCT data set of the 2^nd^ block (t_(27)_ = 2.59, p = 0.008, VCT decrease in 18 out of 28 individuals, MD = −7.85 ± 21.84%, **Figure 6C**).

Similarly to experiment 1, we assessed the HR of cutaneous sensation detection (HR at 0.75mA = 16.07 ± 27.4%, 1mA = 21.43 ± 30.21%, 1.5mA = 50.89 ± 43.29%, mean HR = 29.46 ± 27.36%). We reanalyzed our main outcome parameter by adding mean cutaneous sensation detection HR as a covariate (HRs were z-scored because of non-normal distribution). The main effect of *hf-tACS_sine_* remained significant (F_(3, 78)_ = 4.73, p = 0.004, 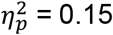). The mean HR of cutaneous sensation did not correlate with the average hf-tACS_sine_-induced VCT modulation (r = −0.12, p = 0.542). In this experiment we additionally tested phosphenes detection (HR_phos_ at 0.75mA = 3.57 ± 8.91%, 1mA = 5.36 ± 12.47%, 1.5mA = 6.25 ± 16.14%, mean HR = 5.06 ± 10.48%). After adding HR_phos_ as covariate (z-scored HR_phos_), the main effect of *hf-tACS_sine_* remained significant (F_(3, 78)_ = 4.69, p =^2^0.005, 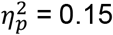). Accordingly, the mean HR of phosphene detection did not correlate with the average hf-tACS_sine_-induced VCT modulation (r = −0.14, p = 0.493).

### 3.3 Comparison of hf-tACS_triangle_, hf-tACS_sine_, and hf-tRNS-induced modulation

First, we tested whether baseline VCT measured during the no tES conditions differed between the experiments (i.e., average baseline VCT in hf-tACS_triangle_, hf-tACS_sine_, and hf-tRNS experiments, see **Figure 7**). Bayesian independent samples t-test revealed that the baseline VCT measured in the no tES condition did not differ between experiments (BF_10_ = 0.21, i.e., moderate evidence for the H_0_).

**Figure 7.**
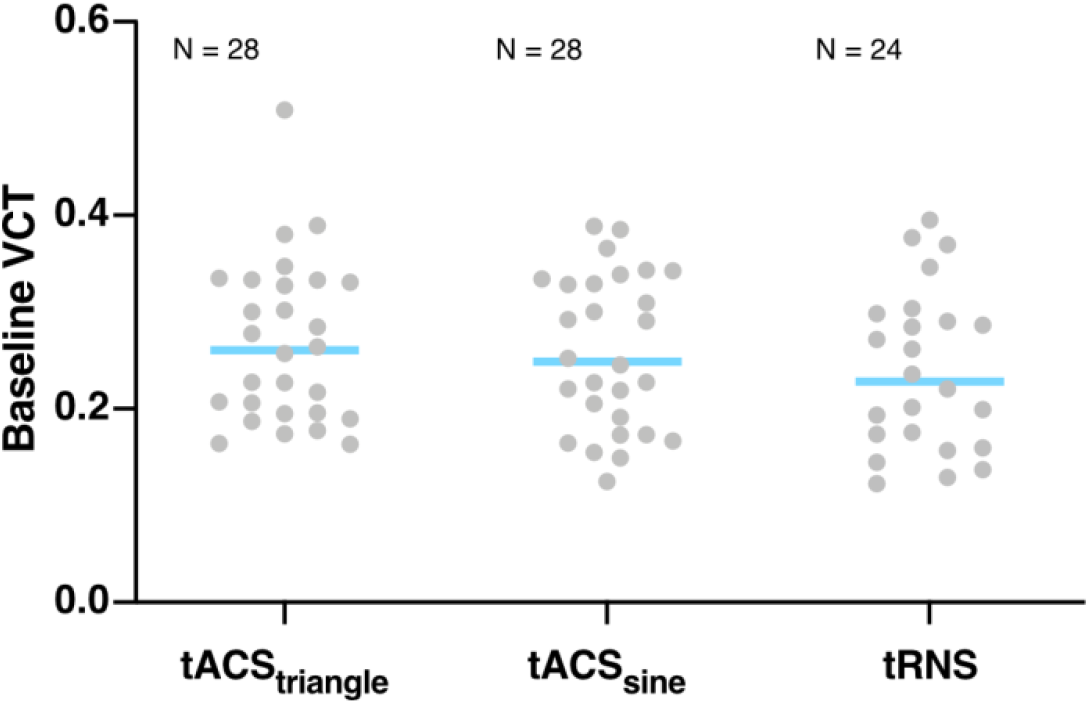
Average baseline VCT measured in the no tES conditions in hf-tACStriangle, hf-tACSsine, hf-tRNS experiments. VCT was assessed for stimuli presented with contrast intensity ranging from 0 to 1. Blue lines indicate mean, gray dots indicate single subject data.

Next, we compared tES-induced modulation effects between experiments (hf-tACS_triangle_, hf-tACS_sine_ and hf-tRNS experiments, see **Figure 8A**). A Bayesian ANOVA revealed that the general tES-induced modulation did not differ between experiments (BF_10_ = 0.11, i.e., moderate evidence for the H_0_), suggesting that all three stimulation types were equally effective in lowering VCT.

**Figure 8.**
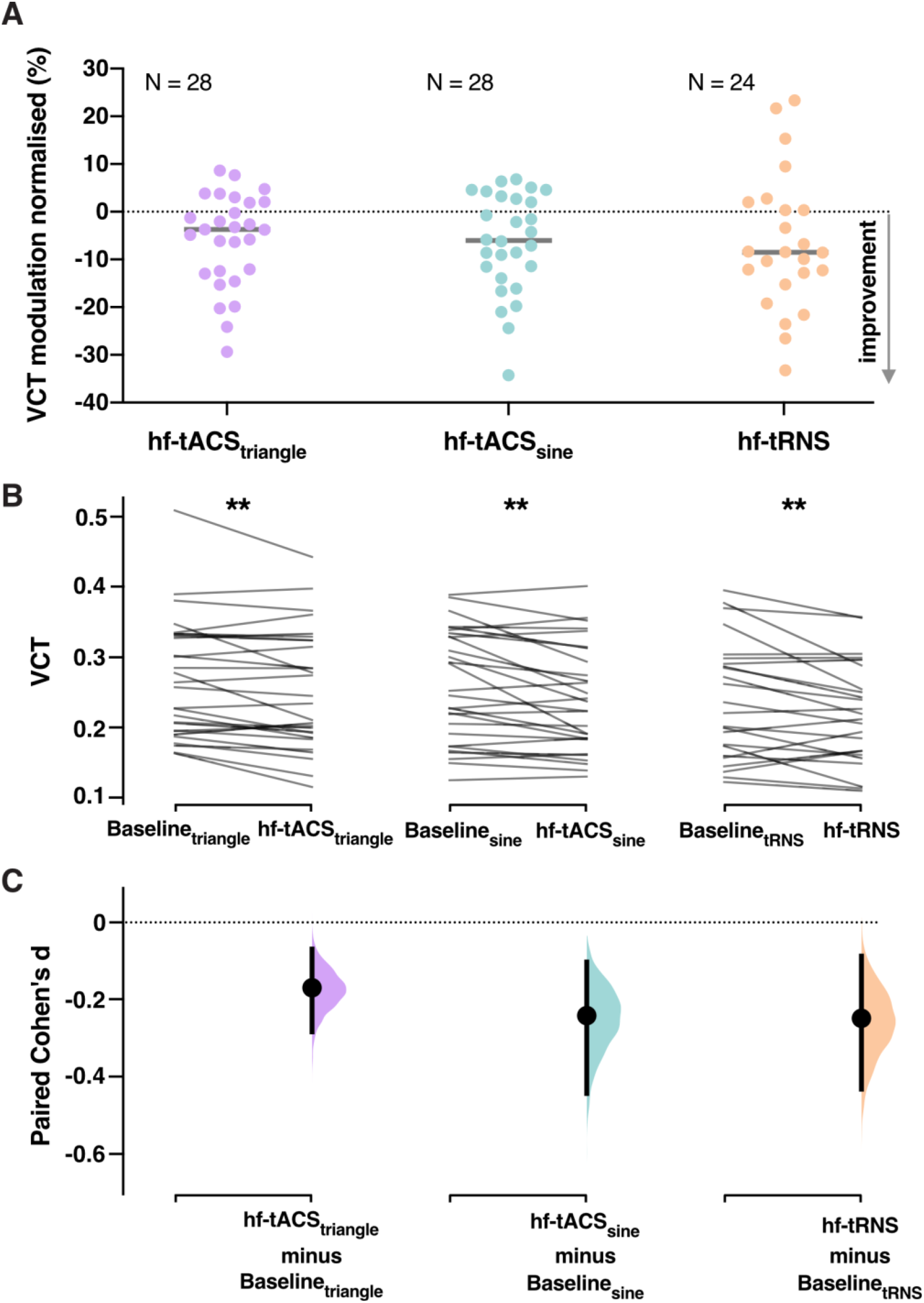
Comparison of hf-tACS_triangle_, hf-tACS_sine_, and hf-tRNS-induced modulation. **A**. VCT modulation induced by hf-tACS_triangle_, hf-tACS_sine_, hf-tRNS. The general modulation of VCT induced by tES was calculated as mean of all active tES conditions from 1st and 2nd blocks normalized to baseline no tES condition in each experiment. Decrease in VCT reflects improvement of visual contrast sensitivity. **B**. Pairs of raw VCT data from hf-tAC_Striangle_, hf-tACS_sine_, hf-tRNS experiments. VCT was assessed for stimuli presented with contrast intensity ranging from 0 to 1. Lines represent each paired set of observations. **C**. The paired Cohen’s d for 3 comparisons shown in the Cumming estimation plot. Each paired mean difference is plotted as a bootstrap sampling distribution. Mean differences are depicted as dots, 95% confidence intervals are indicated by the ends of the vertical error bars. **p < 0.01.

Finally, we assessed the strength of the tES-induced effects on VCT across hf-tACS_triangle_, hf-tACS_sine_ and hf-tRNS experiments defined as paired Cohen’s d bootstrapped sampling distributions (see **Figure 8B-C**). We found comparable (small) effects of significant differences between no tES baseline VCT and averaged VCT in active tES conditions in all experiments using the two-sided permutation t-test [in hf-tACS_triangle_ d = −0.17 (95.0%CI −0.284; −0.0698) p = 0.0034; in hf-tACS_sine_ d = −0.242 (95.0%CI −0.444; −0.103), p = 0.0016; in hf-tRNS d = −0.249 (95.0%CI −0.433; −0.088) p = 0.0092]. The effect sizes and CIs are reported above as: effect size (CI width lower bound; upper bound).

## 4. Discussion

Theoretical modelling shows that adding a deterministic signal instead of stochastic noise can lead to enhanced signal detection. In the present study, we investigated whether stimulation of V1 with a deterministic hf-tACS signal instead of stochastic noise leads to signal enhancement in visual processing. We measured visual contrast sensitivity during hf-tACS_triangle_ and hf-tACS_sine_. On the group level, we found consistent hf-tACS_triangle_- and hf-tACS_sine_-induced decrease in VCT, reflecting enhancement in visual contrast processing during V1 stimulation (**Figure 5A**, **Figure 6A**). The online modulation effects of individually optimized hf-tACS_triangle_ and hf-tACS_sine_ intensities (**Figure 5B**, **Figure 6B**) were replicated on the independent VCT data (**Figure 5C**, **Figure 6C**). Finally, we demonstrated that the effects of non-stochastic stimulation on VCT are comparable to stochastic stimulation of V1 with hf-tRNS (**Figure 8A-C**).

### 4.1 hf-tACS with triangle and sine waveform improve visual sensitivity

Our findings provide the first proof of concept that non-stochastic hf-tACS_triangle_ and hf-tACS_sine_ delivered to V1 can modulate visual contrast sensitivity. Across two experiments we showed that the modulatory effects of hf-tACS on visual sensitivity are not waveform specific, as both hf-tACS_triangle_ and hf-tACS_sine_ induced significant decrease in VCT (**Figure 5A**, **Figure 6A**).

One of the main characteristics of SR-like effects is the optimal intensity of noise, which is required in order to yield the improved performance (Dykman and Mcclintock, 1999; Moss et al., 2004). Here, we did not observe an excessive level of hf-tACS that would be detrimental for visual processing (**Figure 5A**, **Figure 6A**). This is consistent with our predictions that adding high frequency deterministic signal should result in a noise-free output where the detection processing is not disturbed by random stimulation effects.

Similar to other studies investigating resonance-like effects (Potok et al., 2022a; van der Groen and Wenderoth, 2016), our results have revealed large variability among participants in terms of the optimal intensity resulting in the strongest modulation of visual contrast sensitivity (**Figure 5B**, **Figure 6B**). However, consistent with the effects of tRNS-induced online modulation of contrast processing in V1 shown previously (Potok et al., 2022a; van der Groen and Wenderoth, 2016), the effects of individualized hf-tACS intensity were replicated on the independent VCT data set collected within the same experimental session (**Figure 5C**, **Figure 6C**), suggesting consistent beneficial resonance-like influence of hf-tACS on signal enhancement.

We implemented several control measures to test whether the improvement in visual processing was driven by effective stimulation of V1 rather than any unspecific effects of hf-tACS. We applied an anesthetic cream to numb potential stimulation-induced cutaneous sensation on the scalp (Asamoah et al., 2019; van der Plas et al., 2020). While the anesthetic cream numbs the skin and reduces the cutaneous sensations resulting from tACS, it does not eliminate them completely in all individuals. The control cutaneous sensation detection assessment in the current study showed that some participants could accurately detect hf-tACS, and that the mean detection rate was rather low (mean HR = 24.11 ± 27.34% in hf-tACS_triangle_ and mean HR = 29.46 ± 27.36% in hf-tACS_sine_). Cutaneous sensation and phosphenes detection (also very low, mean HR_phos_ = 5.06 ± 10.48%) did not correlate with the average hf-tACS-induced VCT modulation neither in hf-tACS_triangle_, nor hf-tACS_sine_ experiment. Moreover, stimulation effects remained significant in the additional analysis using tactile or phosphene sensation detection during hf-tACS_triangle_ and hf-tACS_sine_ as covariate.

While tACS_sine_ is a well-established and frequently used non-invasive brain stimulation method, hf-tACS_sine_ is less common. The effects of 80Hz tACS_sine_ were sporadically tested in the past using physiological and behavioral paradigms. Ten minutes of 140Hz tACS_sine_ was shown to increase primary motor cortex (M1) excitability as measured by transcranial magnetic stimulation-elicited motor evoked potentials during and for up to 1h after stimulation. Control experiments with sham and 80Hz stimulation did not show any effect, and 250Hz stimulation was less efficient, with a delayed excitability induction and reduced duration (Moliadze et al., 2010). The researchers postulated that the changes in corticospinal excitability result from externally applied high frequency oscillation in the ripple range (140Hz corresponding to middle, 80Hz lower and 250Hz upper border) that interfere with ongoing oscillations and neuronal activity in the brain (Moliadze et al., 2010). We cannot directly translate the effects of hf-tACS_sine_ of M1 to our stimulation of V1. Additionally, the stimulation effects observed in our study are likely reflecting acute modulation of contrast processing, as stimulation was only applied for short intervals (2 s) always interleaved with control (no hf-tACS) condition. Thus, it is possible that even though 80Hz stimulation did not lead to long term effects in cortical excitability it can still affect cortical processes acutely.

In the visual domain, 1.5mA tACS_sine_ was applied to V1 for 15-45min in a study investigating the effect of covert spatial attention on contrast sensitivity and contrast discrimination (Laczó et al., 2012). They found that contrast discrimination thresholds decreased significantly during 60Hz tACS_sine_, but not during 40 and 80Hz stimulation. This previous study used, however, different visual stimuli than the utilized here, i.e., a random dot pattern. Moreover, they used a more complicated behavioral paradigm, where contrast-discrimination thresholds were tested using two attention conditions, i.e., with or without peripheral cue, as the study goal was to explore the influence of attentional processes on visual tasks.

Even though the vast majority of tACS studies to date have used a sinusoidal waveform, an alternating current does not have to be sinusoidal, since it can take any arbitrary waveform such as rectangular wave (Marshall et al., 2006), pulsed (Jaberzadeh et al., 2014), or sawtooth (Dowsett and Herrmann, 2016). Dowsett and Herrmann (2016) investigated the effects of sinusoidal and sawtooth wave tACS on individual endogenous alpha-power enhancement. They observed alpha oscillations enhancement both during and after sawtooth stimulation. The effect seemed to depend on the shape of the sawtooth, as they found that positive, but not negative, ramp sawtooth significantly enhanced alpha power during stimulation relative to sham. They postulated that a sudden, instantaneous change in current might be more effective than a sinusoidal current in increasing the probability of neurons firing. In this regard, Fröhlich and McCormick (2010; Supplementary Material) demonstrated that ramps of increasing voltage with a steeper gradient resulted in increased neural firing in vitro, relative to ramps with a low gradient but reaching the same maximum voltage. This suggests that it is not only the total amount of current but also the rate of change of current can modulate neural firing. Note, that triangle waveform has a faster rate of change of current than the sine wave.

Although we postulate that the effect of hf-tACS on VCT in our study results from resonance-like mechanism, this is not the only potential mechanism. Importantly, the commonly accepted mechanism of action of tACS is that it entrains action potential firing, and thus neural oscillations (Fröhlich et al., 2014). Entrainment effect anticipates a monotonic relationship between the tACS effect and intensity, where increasing stimulation intensity results in greater effects for stimulation waveforms that are tuned to the endogenous oscillation (Thut et al., 2017). The effects of tACS in regard to induced brain oscillations seem to depend on the stimulation duration (Herrmann and Strüber, 2017). Although the entrainment after-effects were observed after tACS delivered for several minutes (for review see Herrmann and Strüber, 2017), short stimulation of 1s did not produce after-effects on amplitude or phase of the electroencephalogram (Strüber et al., 2015). Moreover, in the study investigating the effects of intermittent alpha tACS of either 3 or 8 s, the after-effects were found only for the 8-s condition (Vossen et al., 2015). The authors excluded entrainment as potential underlying mechanism and postulated plasticity-related changes as the responsible mechanism for the observed after-effects. Here, we used very brief stimulation of 2s tACS per trial, a duration seemingly too short to induce the entrainment effects on cortical processing.

Furthermore, it was postulated that a very small amount of applied electric field can bias spike timing or spike probability when a neuron nears the threshold of spike generation (Liu et al., 2018). Accordingly, it was shown that although entrainment effects can arise at field strengths <0.5 mV/mm, physiological effects are more pronounced for higher intensities (around 1mV/mm), according to intracranial recordings in awake nonhuman primates (Johnson et al., 2020). These values are well above the simulated induced electric field in our study (around 0.2 mV/mm, see **Figure 3**). Further studies are required to fully disentangle the underlying neuronal effects of hf-tACS driving the enhancement in visual detection. To exclude the influence of entrainment on VCT modulation a jittered hf-tACS protocol could be employed. A paradigm using stimulation of jittered flickering light, where instead of a rhythmic flicker, inter stimulus intervals of the square wave were jittered with a maximum of ± 60%, was shown to fail in inducing rhythmic brain response (Notbohm et al., 2016). If a jittered hf-tACS of V1 would still influence contrast sensitivity we could assume the non-entrainment origin of the effect.

### 4.2 Comparison of hf-tACS_triangle_, hf-tACS_sine_, and hf-tRNS

In the phenomenon of SR, random noise added to a non-linear system can increase its responsiveness towards weak subthreshold stimuli. One aim in the present study was to explore whether a deterministic and periodic signal can substitute stochastic noise and still lead to response enhancement in a threshold-based stochastic resonator. Non-stochastic characteristics of high-frequency deterministic signal might offer a noise-free output, thus additionally increasing SNR. We proposed the following testable hypotheses (i) hf-tACS_triangle_ will have a larger resonance-like effect compared to hf-tRNS, (ii) hf-tACS_sine_ will have less effect than hf-tACS_triangle_, due to the loss of waveform linearity. We found enhancement effects of both hf-tACS_triangle_ vs hf-tACS_sine_ (**Figure 5A**, **Figure 6A**), however to test whether these effects are indeed superior to stochastic stimulation, we directly compared the VCT modulation induced by hf-tACS_triangle_, hf-tACS_sine_ and hf-tRNS (**Figure 8**). The baseline contrast sensitivity between the compared experiments was not different (**Figure 7**). Counter to our hypothesis, the noise-free hf-tACS did not result in stronger contrast sensitivity enhancement, as average VCT modulation did not differ between the three stimulation conditions, as confirmed by Bayesian analysis (**Figure 8A**). Accordingly, the effects sizes of all three stimulation types were comparable (**Figure 8B-C**). Therefore, we showed that both non-stochastic and stochastic high-frequency stimulations were equally effective in inducing resonance-like effects.

### 4.3 Conclusions

The present study demonstrates the first evidence of the resonance-like neural signal enhancement without stochastic noise component. We showed that ‘non-stochastic’ hf-tACS and ‘stochastic’ hf-tRNS are equally effective in enhancing visual contrast detection. In the range of commonly used intensities of tES to induce SR, hf-tACS did not result in detrimental effects related to excessive interference signal, thus providing increased SNR in all tested intensities. These findings shed a new light on the effects induced by both hf-tACS and hf-tRNS, and their underlying mechanisms.

## 13. ACKNOWLEDGEMENTS

We thank all the participants for their time and effort.

